# Changes in slow oscillations and sleep spindles by auditory stimulation positively correlate with memory consolidation in children with epilepsy and controls

**DOI:** 10.1101/2025.05.12.653459

**Authors:** Hunki Kwon, Dhinakaran M. Chinappen, Anirudh Wodeyar, Elizabeth Kinard, Skyler Goodman, Wen Shi, Bryan S. Baxter, Dara S. Manoach, Mark A. Kramer, Catherine J. Chu

**Affiliations:** Department of Neurology, Massachusetts General Hospital, Boston, Massachusetts, USA; Harvard Medical School, Boston, Massachusetts, USA; Department of Psychiatry, Massachusetts General Hospital, Boston, Massachusetts, USA; Athinoula A. Martinos Center for Biomedical Imaging, Charlestown, MA; Department of Mathematics and Statistics, Boston University, Boston, Massachusetts, USA; Center for Systems Neuroscience, Boston University, Boston, Massachusetts, USA; Department of Advanced Computing Sciences, Maastricht University, Maastricht, The Netherlands

## Abstract

**Background:** Sleep-dependent memory consolidation is supported by sleep spindles during stages 2 and 3 non-rapid eye movement sleep. Sleep spindles and sleep-dependent memory consolidation are both decreased in Rolandic epilepsy (RE). Non-invasive auditory stimulation evokes SOs and SO-spindle complexes in healthy adults but the impact on memory consolidation has been inconsistent.

**Objective:** We investigated the effects of auditory stimulation during sleep on SOs, SO-spindle complexes, and sleep-dependent memory consolidation in children with RE and controls.

**Methods:** A prospective cross-over study was conducted in children with RE and control. Children completed two nap visits with auditory or sham stimulation. SOs and SO-spindle complexes rates were measured offline using validated detectors. Sleep-dependent memory consolidation was assessed using the motor sequence typing task.

**Results:** Auditory stimulation evoked SOs and SO-spindle complexes broadly with maximal effect over frontal electrodes. Compared to sham, stimulation delivered during background activity evoked SOs (29.8% increase, p<0.001) and SO-spindle complexes (16.8% increase, p<0.001) and stimulations delivered near the peak of an ongoing SO upstate maximally evoked SOs (51.3% increase, p<0.001) and SO-spindle complexes (32.3% increase, p<0.001). Changes in frontal SO (1.9% improvement per increase in SO/min; p<0.001) and SO-spindle complexes (9.5% improvement per increase in SO-spindle/min) event rates due to auditory stimulation positively predicted changes in sleep-dependent memory consolidation.

**Conclusion:** Auditory stimulation reliably modulates sleep oscillations when delivered on background activity and during the upstate of SOs. As increased event rates improve memory consolidation, stimulation paradigms to increase SO and SO-spindle complex rates are required to enhance memory.

## 1. Introduction

Sleep is critical for memory consolidation -- the integration of newly acquired information into long-term storage^1^. Slow oscillations (SOs), large low frequency brain rhythms (0.5–2 Hz) during deep non-rapid eye movement (NREM) sleep synchronize thalamocortical spindles (bursts of 10-14 Hz oscillations^2^) and hippocampal sharp wave ripples (80-120 Hz), facilitating communication between the hippocampus and neocortex^1^. The depolarizing SO upstate facilitates the initiation of sleep spindles by GABAergic neurons in the thalamic reticular nucleus^3,4^, leading to an increased occurrence of spindles observed in the cortex and thalamus compared to baseline periods^5-7^. Sleep spindles may facilitate synaptic plasticity by regulating dendritic calcium shifts^8^ and coordinating hippocampal sharp wave ripples that drive neuronal replay and contribute to memory consolidation^9,10^. The functional coupling of SOs, spindles, and hippocampal sharp wave ripples during NREM sleep supports synaptic potentiation and interregional communication, essential for memory consolidation^1,11,12^.

Rolandic epilepsy (RE), also known as self-limited epilepsy with centrotemporal spikes (SeLECTS), is the most common focal developmental epilepsy in childhood, accounting for 8–23% of childhood epilepsy^13^. RE is characterized by sleep-activated spikes in the inferior Rolandic cortex and varying degrees of cognitive deficits during school-age years^13^, The most prominent cognitive symptom in RE is memory impariment^14^, and includes deficits in sleep-dependent memory consolidation^15^. In RE, cortical regions with epileptiform spikes also have a paucity of sleep spindles^16^ and both sleep-dependent memory and IQ correlate with spindle rate^15,16^.

Work in the last decade has demonstrated that auditory stimulation phase-targeted to the upstate of endogenous SOs during NREM sleep can induce a subsequent SO and associated spindle activity^17-23^. This increase in SOs and spindles was initially reported to improve declarative memory consolidation in healthy adults^17,18,24^ and in typically developing children^23^. However, subsequent studies found that auditory stimulation reliably evokes SOs and SO-sleep spindle complexes, but does not reliably improve declarative or procedural memory consolidation in healthy adults^20,22,25^ or children^23^. Importantly, although prior work suggests that the rates of SOs, sleep spindles, and SO-spindle complexes best predict memory consolidation^15,26-29^, most studies evaluating the impact of auditory stimulation on memory consolidation did not evaluate the overall rate of these oscillations. When applied in epilepsy, one prior study found that auditory stimulation during NREM sleep suppressed spike activity in children with RE^30^, however, the impact on sleep oscillations supporting memory and consequent sleep-dependent memory consolidation was not evaluated^30,31^.

In this study, we hypothesized that auditory stimulation during stages 2 and 3 NREM sleep would evoke SO and SO-spindle complexes in children with RE and controls, compared to sham stimulation delivered at the same phase. We also hypothesized that changes in SO and SO-spindle complex rates due to auditory stimulation would predict changes in sleep-dependent memory consolidation. To test these hypotheses, we performed a prospective cross-over study and evaluated the rates of SOs and SO-spindle complexes and sleep-dependent memory consolidation during a nap with randomly time auditory stimulation compared to a nap with sham stimulation. Understanding how auditory stimulation impacts sleep oscillations and memory consolidation will enable translation of this non-invasive, scalable, neuromodulatory approach to enhance cognitive function in children impacted by epilepsy and other conditions that impact memory.

## 2. Materials and Methods

### Subjects

We prospectively recruited children with Rolandic epilepsy (RE) and control children to participate. All subjects that enrolled in the study reported in ^15^ were invited to participate in this study. Diagnosis of RE was confirmed by a pediatric epileptologist (C.J.C.) based on International League Against Epilepsy criteria, requiring a history of at least one focal motor or generalized seizure and EEG evidence of sleep-activated centrotemporal spikes^32,33^. Children with attention disorders or mild learning difficulties were included as these are common symptoms in RE^13,14^. Informed consent was obtained from all participants, and the study was approved by the Massachusetts General Hospital Institutional Review Board.

### Study design

The study consisted of two visits to the Athinoula A. Martinos Center for Biomedical Imaging. For each visit, subjects arrived at approximately 10:00 AM and began with motor sequence task training (**Figure 1A**). This was followed by a nap opportunity lasting approximately 90 minutes, scheduled between 1:00 PM and 3:00 PM. After the nap, subjects completed the motor sequence task testing. Each visit was assigned to either no stimulation or auditory stimulation (see Auditory Stimulation). In each case, the EEG was monitored continuously during the nap opportunity and stimulation was delivered throughout stage 2 and stage 3 NREM sleep.

**Figure 1.**
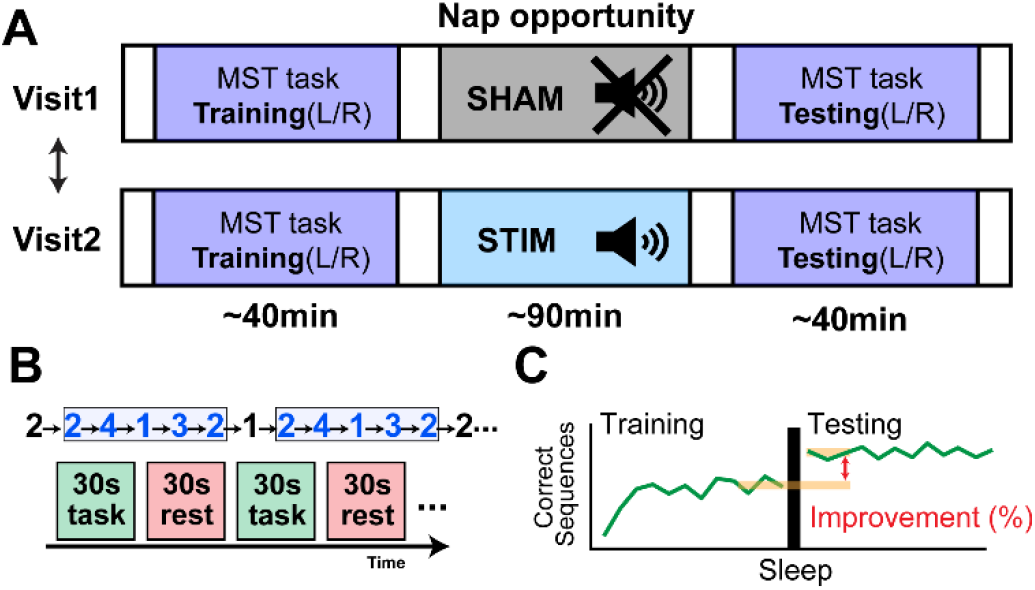
Experimental overview. **A)** One visit included auditory stimulation with randomly timed 50 ms bursts of pink noise (STIM) during Stages 2 and 3 NREM sleep, and one visit included no auditory stimulation (SHAM). **B)** Subjects performed the finger tapping motor sequence task with their left and right hand separated by a 10-minute break. **C)** Sleep-dependent memory consolidation is the percentage difference between the number of correct sequences during the last three trials of training and the first three trials of testing.

### Motor sequence task

To measure sleep-dependent memory consolidation, subjects completed the finger tapping motor sequence task (MST) as in ^15^. This involved typing with the left hand a 5-element sequence on a labeled number pad (e.g., 2-4-1-3-2, **Figure 1B**) as quickly and accurately as possible for twelve 30 second trials, each separated by a 30-second rest interval^34,35^. The sequence was displayed during trials to minimize working memory load. After a 10-minute break, subjects performed the task with their right hand using a different sequence. To exclude performance outliers across trials, for each subject and each hand, we fit an exponential model to the learning curve during training, and included a constant offset to the exponential model during the post-sleep testing, as in ^15,36^. If performance on a trial was less than expected (defined as more than two standard deviations below the model fit), it was excluded as an outlier. Outliers could be due to misplaced fingers or task interruption, for example. Sleep-dependent memory consolidation was calculated as the percentage difference between the mean performance of the last three trials before sleep and the first three trials after awakening and measured separately for the left- and right-hand MST (**Figure 1C**), as done previously^15^.

### EEG recordings

EEG recordings were acquired using 70-channel high density EEG caps (Easycap, Vectorview, Elekta-Neuromag, Helsinki, Finland) at 2035 Hz or 2000 Hz sampling rates. Data were downsampled to 407 Hz or 400 Hz for analysis. Electrode impedances were maintained below 10 kΩ. Electrode positions were digitized using a 3D digitizer (Fastrak; Polhemus Inc., Colchester, VA). A board-certified neurophysiologist visually inspected the data to identify channels with poor recording quality and sleep staged following standard procedures^37^.

### Auditory stimulation

Auditory stimulation was delivered using Psychtoolbox^38^ in MATLAB (The MathWorks, R2021b, Natick, MA), while EEG data were monitored in real-time by a board-certified neurophysiologist (C.J.C.). When subjects entered stages 2 and 3 NREM sleep, 50 ms bursts of pink noise were randomly presented through in-ear headphones, with an interstimulus interval ranging from 2 to 30 seconds. The volume of auditory stimulation was set based on subject feedback prior to the nap opportunity, adapted for comfort without disturbing sleep, within a range of 32 to 35 dB. The ambient noise level in the room was ∼30dB. To prevent sleep disruptions, stimulations were paused manually if arousals or awakenings occurred, and distractions in the recording environment (e.g., noise in the recording room) were minimized.

### Offline slow oscillation detector

SOs were identified offline after referencing sleep-staged artifact-free EEG data to the nasion. Stages 2 and 3 NREM EEG data were bandpass filtered (0.5–4 Hz) and SOs detected if two consecutive positive-to-negative zero crossings occurred within 0.5–2 seconds and the negative peak during this interval was less than or equal to -40 μV, adapted from ^39^.

### Offline sleep spindle detector

For spindle detection, stages 2 and 3 NREM EEG data were re-referenced to an average reference. To detect sleep spindles, we used an automated spindle detector, developed to perform well in the setting of sharp events in EEG, such as pediatric vertex waves or epileptiform spikes^2,16^. Spindles were required to last at least 0.5 seconds^40,41^ and spindles detected within 1 second of each other were concatenated^16,40^.

### Evoked SO and SO-spindle complex percentage calculation

Randomly timed auditory stimulations were delivered during NREM stages 2 and 3 sleep. The percentage of stimulations that evoked SOs was calculated as the proportion of SOs occurring within 1 second following auditory stimulation divided by the total number of stimulations multiplied by 100. The percentage of stimulations that evoked SO-spindle complexes was determined as the number of spindles occurring within 1 second after the negative peak of an evoked SOs following auditory stimulation divided by the total number of stimulations multiplied by 100. These percentages were calculated separately for stimuli delivered during baseline activity and for stimuli delivered during an endogenous SO (i.e., a spontaneous SO not evoked by auditory stimulation). For stimulations delivered during endogenous SOs, percentages were calculated across 12 phase bins, each representing 30° increments of the endogenous SO phase. To estimate the endogenous SO phase, the EEG data were bandpass filtered (0.5–2 Hz) and the smoothed phase was estimated after correcting for waveform distortions caused by narrowband filtering^42^. This approach enabled us to evaluate the proportion of auditory stimulations that evoked a SO and/or SO-spindle complex relative to the endogenous SO phase at the time of auditory stimulation. Percentages were then compared to sham stimulations delivered during baseline activity or at the same phase of an endogenous SO.

### Statistical analyses

To identify significant intervals of times where the EEG signals, spindle detections, evoked SO, or evoked SO-spindle complex percentages differed between auditory stimulation and sham stimulation, temporally contiguous cluster-based permutation tests were performed^2,43^. A two-sided paired t-test was first used to identify time points (or phase bins) that exhibited significant differences between stimulation conditions across subjects. Time points (or phase bins) with a p-value below the critical alpha levels (p < 0.05 and p< 0.01) were clustered if the p-values were temporally adjacent within 50 ms (or adjacent based on phase bins). Temporally contiguous cluster-level statistics were determined by summing the absolute t-values within each cluster. To determine which temporally contiguous cluster-level statistics were unlikely to occur by chance, condition order was randomly shuffled within each subject, and the largest temporally contiguous cluster-level statistic was measured from the resampled data using the same method as for the unpermuted data. Temporally contiguous clusters from the unpermuted data were considered significant if their statistics exceeded the top 5% and 1%of statistics computed from 5000 resamplings of the data. Identical procedures were used to detect differences in phases of the endogenous SOs that evoked spindle detections, evoked SO detections, and evoked SO-spindle complex events. Identical procedures were also used to identify spatial clusters of electrodes where the evoked SO or SO-spindle complex percentage differed between auditory stimulation and sham stimulation, except that a spatially contiguous cluster-based permutation test was performed instead of a temporally contiguous cluster-based permutation test^2,43^. For this approach, channels with a p-value below the critical alpha level were clustered if they were spatially adjacent. For visualization, topographic maps of spatially contiguous significant clusters were interpolated using the Fieldtrip toolbox (http://www.ru.nl/neuroimaging/fieldtrip)^44^.

To test for a relationship between event rates (SO or SO-spindle complex rates) and sleep-dependent memory improvement in the combined sham and auditory stimulation groups, we estimated a linear mixed-effects model with sleep-dependent memory consolidation as the dependent variable, event rate, age, sex, and visit order as predictors, and a subject-specific intercept to account for four observations per subject (left and right hemisphere; sham and auditory stimulation). We assumed that event rates were linked to memory improvement with the contralateral hand^45^.

To examine how changes in event rates (SO or SO-spindle complex rates) induced by auditory stimulation impacted sleep-dependent memory consolidation compared to sham stimulation, we estimated a linear mixed-effects model with the change in sleep dependent memory consolidation with auditory stimulation (compared to sham) as the dependent variable and the change in event rate with auditory stimulation (compared to sham), age, sex, and visit order as predictors, and a subject-specific intercept to account for multiple observations per subject.

### Data availability

Raw data were generated at Massachusetts General Hospital and the Athinoula A. Martinos Center for Biomedical Imaging. Derived data supporting the findings of this study are available from the corresponding author on request. The spindle detection method is available at https://github.com/Mark-Kramer/Spindle-Detector-Method.

## 3. Results

### Subject characteristics

11 children with RE and 9 controls were enrolled. 3 children with RE and 2 controls did not sleep during the nap opportunity and were excluded. In total, 8 children with RE (9.5-17.7 years, 6F) and 7 control children (9.3-16.2 years, 1F) were included. There were no significant difference in age (p=0.55, two-sample t-test) or duration between visits (p=0.12, two-sample t-test) between groups, however more RE subjects were female compared to controls (p=0.02, Chi-square test). All participants were right-handed. Sleep opportunity, stages 2 and 3 NREM sleep duration, and the number of stimulations were not significantly different between sham and stimulation conditions in either group (all p>0.17, paired t-tests). More RE subjects had auditory stimulation during their first visit (5/8) and more control subjects had sham stimulation during their first visit (6/7). Subject characteristics are provided in **Table 1**.

**Table 1.** Subject characteristics. ^a^ Two-sample t-test; ^b^ Chi-square test; ^c^ Paired t-test

|  | RE | Control | p value |
| --- | --- | --- | --- |
| Sample size | 8 | 7 | - |
| Age (years) | 13.3 (9.5-17.7) | 14.2 (9.3-16.2) | 0.55<sup>a</sup> |
| Female sex (%) | 75.0 | 14.3 | 0.02<sup>b</sup> |
| Right handedness (%) | 100 | 100 | - |
| Duration between visits (days) | 11.1 (1-21) | 24.9 (8-71) | 0.12<sup>a</sup> |
| RE |  |  | p value |
| Condition | SHAM | STIM |  |
| Visit 1 | 3 | 5 |  |
| Sleep opportunity (min) | 97.4 (90.0-145.0) | 94.4 (82.0-114.0) | 0.70<sup>c</sup> |
| Sleep duration (N2+N3, min) | 26.5 (3.4-68.9) | 40.9 (9.4-59.9) | 0.17<sup>c</sup> |
| Number of stimulations (n) | 224.8 (26.0-453.0) | 318.9 (64.0-701.0) | 0.32<sup>c</sup> |
| Control |  |  | p value |
| Condition | SHAM | STIM |  |
| Visit 1 | 6 | 1 |  |
| Sleep opportunity (min) | 91.6 (90.0-97.0) | 97.7 (90.0-125.0) | 0.24<sup>c</sup> |
| Sleep duration (N2+N3, min) | 37.0 (8.4-65.3) | 39.3 (17.4-58.5) | 0.83<sup>c</sup> |
| Number of stimulations (n) | 297.4 (61.0-474.0) | 272.4 (122.0-398.0) | 0.75<sup>c</sup> |

### Auditory stimulation during sleep evokes SOs and sleep spindles

Considering the population results from all subjects, regardless of group, auditory stimulation evoked a slow oscillation compared to sham stimulation, indicated by a large triphasic wave with an initial positive deflection near 150 ms, followed by a negative deflection between 300-500 ms, and a second positive deflection near 900 ms (p<0.05, cluster-based statistic, **Figure 2A**). Auditory stimulation tended to reduce spindle activity at the time of stimulation (near 0 ms) followed by a significant increase in sleep spindle detections beginning near 800 ms post-stimulation compared to sham stimulation (p<0.05, cluster-based statistic, **Figure 2B**), temporally coinciding with the second upstate of the induced slow oscillation.

**Figure 2.**
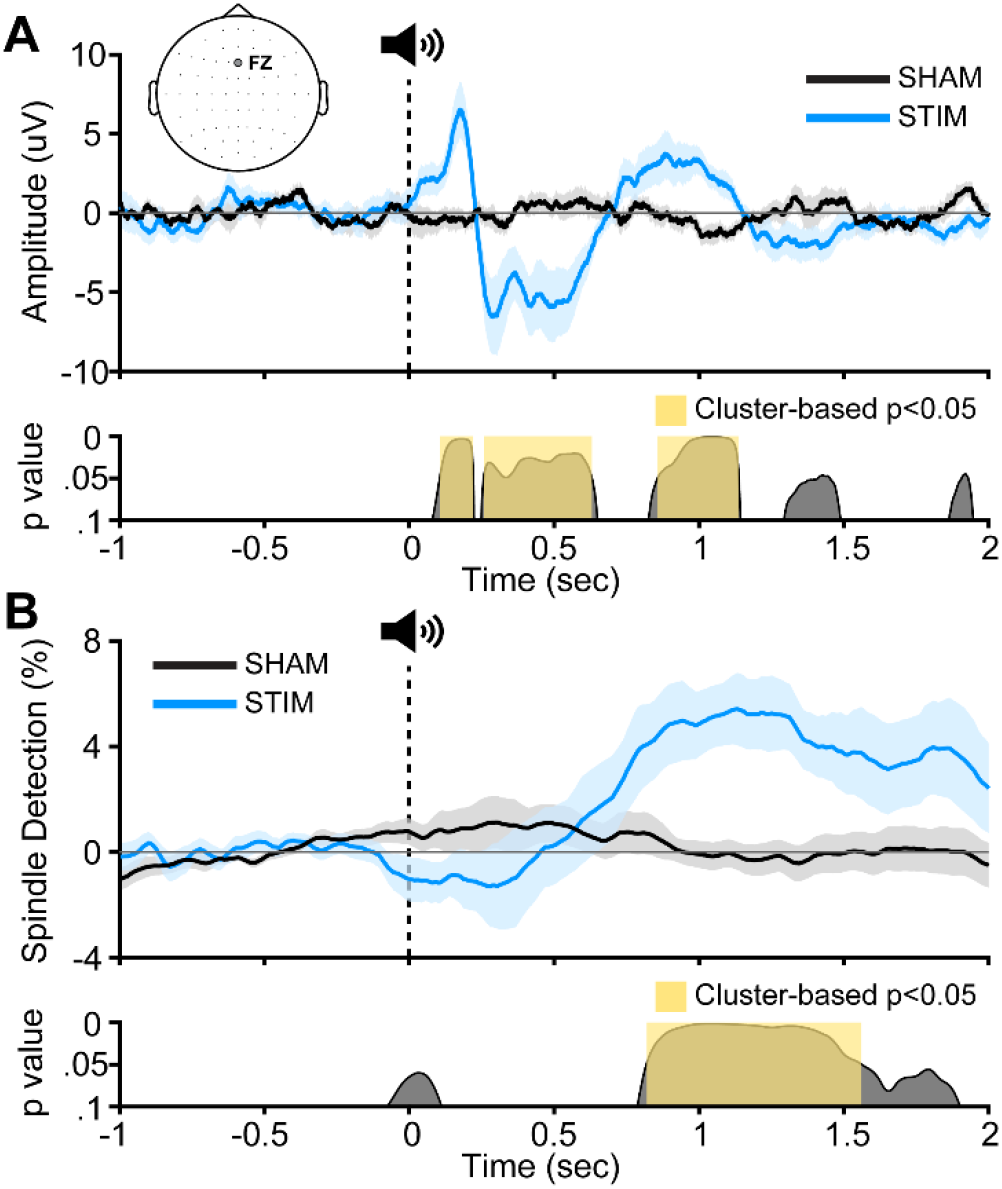
Auditory stimulation evokes SOs and spindles. **A)** Auditory stimulations (averaged across all subjects ± standard area of the mean, blue line) evoke SOs. **B)** Following an initial trend toward reduction in spindles at the time of stimulation, auditory stimulations evoke sleep spindles concurrent with the evoked SO compared to s stimulations (averaged across all subjects, black line). Positive is plotted upward, opposite to EEG convention.

### Topological distribution of evoked SOs and SO-spindle complexes

To examine the spatial distribution of evoked SOs, we evaluated the percentages of evoked SO detected averaged separately for each electrode and compared between stimulation and sham conditions. Stimulations evoked SOs broadly across nearly all scalp electrodes compared to sham stimulation, with maximal effects observed over the frontal, central, and parietal regions near the vertex (p<0.01, cluster-based statistic; **Figure 3A**).

**Figure 3.**
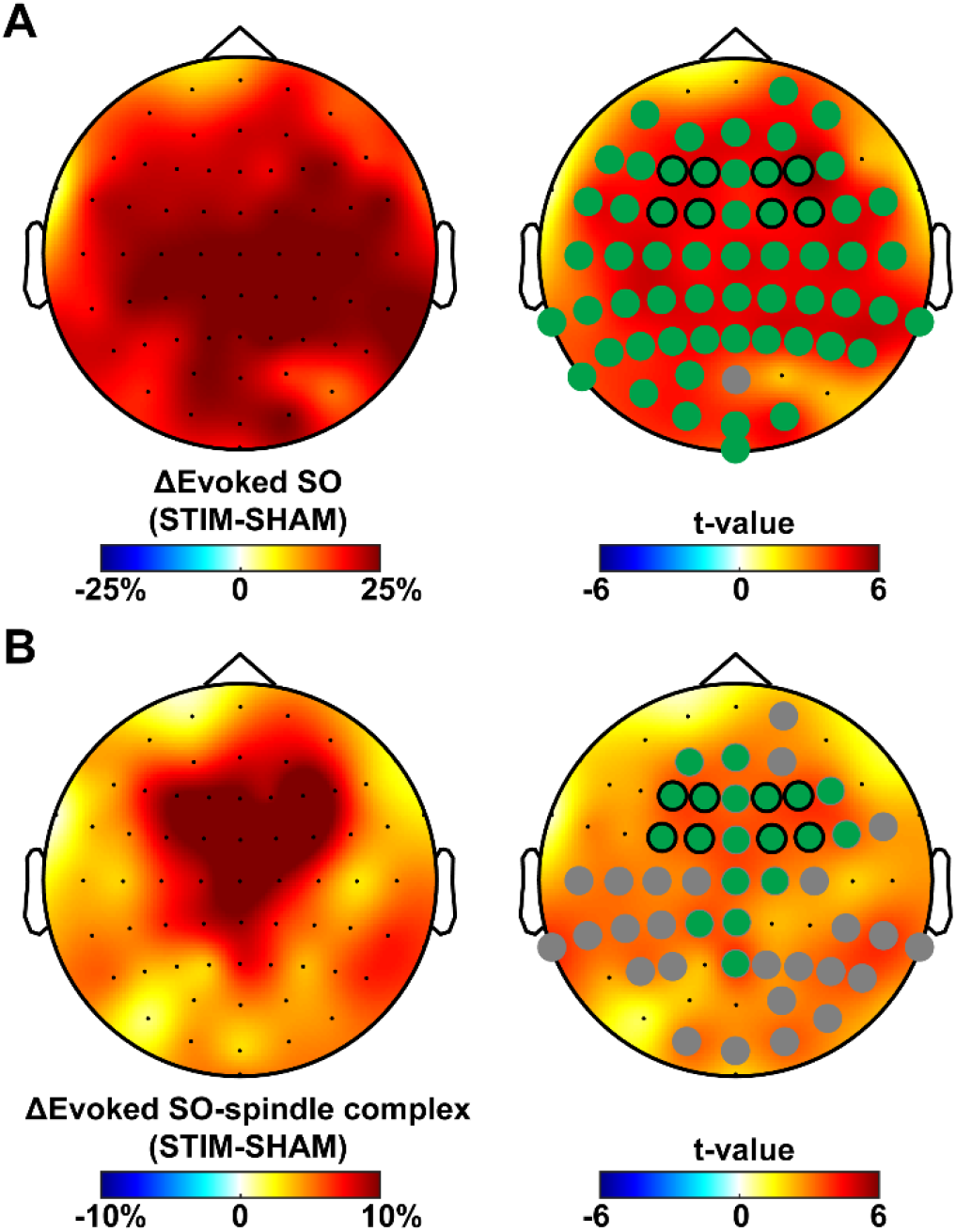
Topological distributions of evoked SO and SO-spindle complexes from auditory stimulation. The distribution of **A)** evoked SO and **B)** evoked SO-spindle complex when auditory stimulation is delivered compared to sham stimulation. Circles indicate electrodes with significantly different event proportions between stimulation and sham conditions (grey: p < 0.05; green: p<0.01, cluster-based statistic). Black-outlined circles indicate symmetric electrodes used to evaluate the impact of auditory stimulation on memory consolidation.

Following a similar analysis approach, the percentages of evoked SO-spindle complexes were averaged separately for each electrode and compared between stimulation and sham conditions. Stimulations evoked SO-spindle complexes broadly (p<0.05, gray **Figure 3B**) but were most prominent over the frontal electrodes near the vertex compared to sham stimulation (p<0.01, cluster-based statistic; green, **Figure 3B**).

These findings indicate that this non-invasive approach can be used to evoke SOs and SO-spindle complexes broadly over the cortex but that the effects are maximal over frontal regions near the midline.

### Auditory stimulation delivered during an endogenous SO upstate preferentially evokes SOs and SO-spindle complexes

To examine the impact of the timing of auditory stimulation relative to endogenous brain activity to evoke SO and SO-spindle complexes, we evaluated the percentage of auditory that evoked a SO delivered in the presence or absence of an endogenous SO compared to sham stimulations delivered on a comparable background (**Figure 4A**). For this analysis, we focused on FZ given the strong effect size observed in topological analysis (**Figure 3**). When auditory stimulations were delivered in the absence of an endogenous SO, an evoked SO occurred 51.0% of the time (Control: 45.4%; RE: 55.9%) and an evoked SO-spindle complex occurred 26.5% of the time (Control: 23.6%; RE: 29.0%). Compared to sham stimulation, the proportion of evoked SOs with auditory stimulation increased by 29.8% (p<0.001); an increase was observed in each group (Control: 21.1% increase, p=0.009; RE: 37.4% increase, p<0.001, two-sided paired t-test; **Figure 4B**). Compared to sham stimulation, the proportion of evoked SO-spindle complexes increased by 16.8% (p<0.001); this was also observed in each group (Control: 13.0% increase, p=0.02; RE: 20.1% increase, p=0.004, two-sided paired t-test; **Figure 4C**).

**Figure 4.**
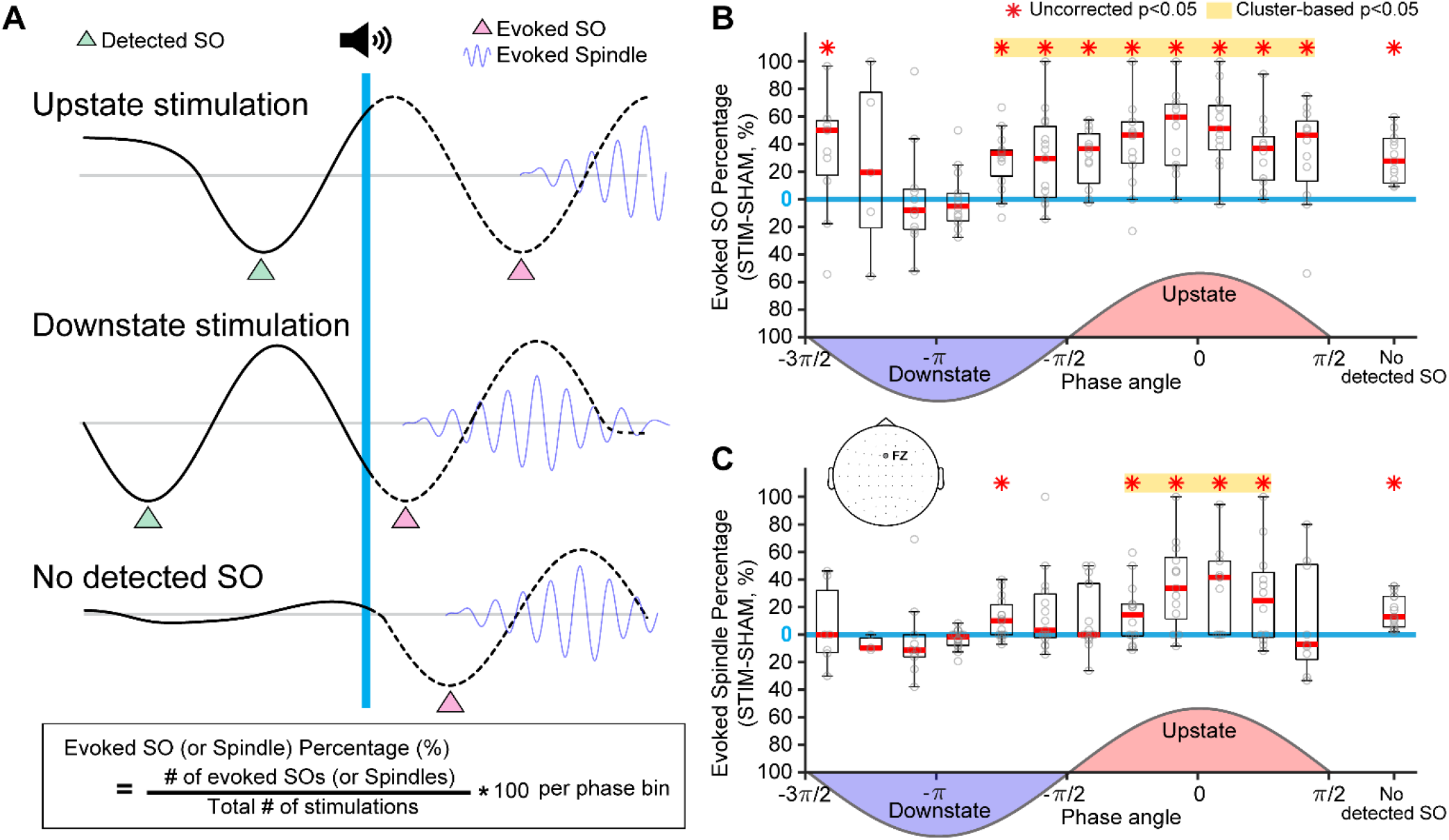
Impact of endogenous activity on stimulation efficacy. **A)** When an endogenous SO is detected (green triangle), the phase is estimated at the time of stimulation (blue vertical line). Stimulation may occur during the endogenous SO upstate (top row) or downstate (middle row). Stimulation may also occur in the absence of an endogenous SO (bottom row). In all cases, an evoked SO (pink triangle) and evoked spindle (blue curve) may occur after auditory stimulation. Evoked responses are compared to sham stimulation delivered on the same background. **B)** Auditory stimulation evokes a higher percentage of SOs and **C)** a higher percentage of SO-spindle complexes when delivered in the upstate of an endogenous SO compared to SHAM. The blue horizontal line indicates the null hypothesis of no difference in evoked SOs and spindles between STIM and SHAM.

When auditory stimulations were delivered at the time of endogenous SO, we evaluated the percentage of auditory stimulations that evoked a SO or SO-spindle complex compared to sham stimulations delivered at the same phase of an endogenous SO. Overall, when stimulations were delivered during the upstate of an endogenous SO, an evoked SO occurred 58.3% of the time (Control: 57.7%; RE: 59.0%) and an evoked SO-spindle complex occurred 32.1% of the time (Control: 32.0%; RE: 32.2%). Compared to sham stimulation, when auditory stimulations were delivered during the upstate of an endogenous SO, the proportion of evoked SOs was increased by 35.7% (p<0.001); this was observed in each group (Control: 30.6% increase, p=0.002; RE: 40.2% increase, p<0.001, two-sided paired t-test; **Figure 4B**). Similarly, when auditory stimulation was delivered during the upstate of an endogenous SO, the proportion of evoked SO-spindle complexes was increased by 23.5% (p=0.002) compared to sham stimulation; this was observed in each group (Control: 21.2% increase, p=0.02; RE: 25.4% increase, p=0.05, two-sided paired t-test; **Figure 4C**). SOs were maximally increased (51.3% increase, p<0.001 compared to sham) when auditory stimulation was delivered at the peak of an endogenous SO upstate (±30 degrees; Control: 45.1% increase, p=0.004; RE: 58.6% increase, p<0.001, two-sided paired t-test). Similarly, SO-spindle complexes were maximally increased (32.3% increase, p<0.001 compared to sham) when auditory stimulation was delivered near the peak of an endogenous SO upstate (±30 degrees; Control: 25.1% increase, p=0.03; RE: 40.7% increase, p=0.01, two-sided paired t-test). In contrast, there was no difference detected in the percentage of SOs and SO-spindle complexes evoked when auditory stimulations were delivered during the downstate of an endogenous SO compared to similarly timed sham stimulation (all p>0.49; **Figure 4B-C**). These findings indicate that auditory stimulation delivered during an upstate of a SO maximally evokes SOs and SO-spindle complexes, but also evokes SO and SO-spindle complexes at all other times except when delivered during a SO downstate.

### Stimulation induced changes in frontal SO and SO-spindle complexes correlate positively with changes in memory consolidation

To evaluate the impact of auditory stimulation on memory consolidation, we evaluated homotopic frontal electrode pairs where the effect size of auditory stimulation was maximal (black outlined circles, **Figure 3B**). Across subjects and visits, both SO rate (p<0.001, mean MST improvement of 0.8%, for each unit increase in SO per minute) and SO-spindle complex rate (p=0.005, mean MST improvement of 5.1%, for each unit increase in SO-spindle complex per minute) positively correlated with memory consolidation (**Figure 5B**). We note that auditory stimulation was not expected to increase event rates since it was delivered randomly. Thus, to evaluate whether changes to neurophysiology during auditory stimulation impact memory, we compared the change in SO rate or SO-spindle complex rate between auditory and sham stimulation visits to the change in sleep-dependent memory consolidation between visits for each subject. We found a positive correlation between the change in SO rate and sleep-dependent memory consolidation (p<0.001, mean MST improvement of 1.9% for each unit increase in SO per minute; **Figure 5C)**. We also found a positive correlation between the change in SO-spindle complex rate and change in sleep-dependent memory consolidation (p=0.007, mean MST improvement of 9.5% for each unit increase in SO-spindle complex per minute).

**Figure 5.**
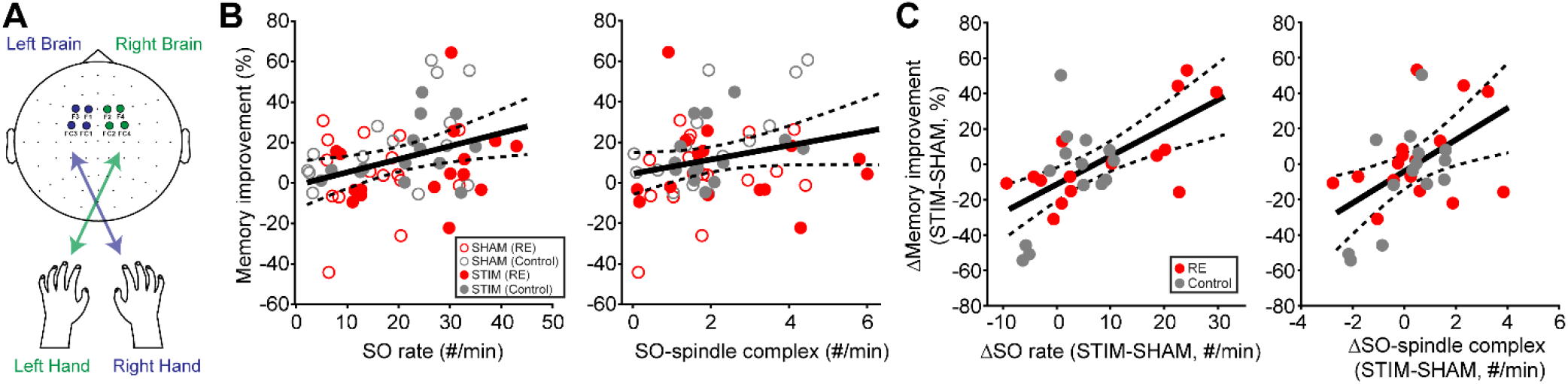
**A)** Changes in MST improvements are compared to changes in frontal sleep event rates between SHAM and STIM visits. Sleep features in the contralateral hemisphere are used to predict motor improvements. **B)** Across all subjects and stimulation conditions, SO rate and SO-spindle coupling rates are positively correlated with sleep-dependent memory improvement **C)** Across all subjects, changes in SO rate and SO-spindle complex rate during auditory stimulation positively correlated with changes in sleep-dependent memory improvement. Black (dashed) curves indicate estimated model fit (95% confidence interval).

Given the small size of our cohorts, we lacked power to evaluate each group separately. However, the change in SO rate was significantly associated with the change in sleep-dependent memory consolidation in RE (p=0.009, mean MST improvement of 2.0% for each unit increase in SO per minute) and the effect size was positive but did not achieve significance in controls (effect size 2.4; p=0.1). Similarly, the change in SO-spindle complex rate was significantly associated with the change in sleep-dependent memory consolidation in controls (p<0.001, mean MST improvement of 24.3% for each unit increase in SO-spindle complex per minute) but was not significant in the RE alone (effect size 0.7, p=0.9). Taken together, these results demonstrate that changes in SO and SO-spindle complex event rates caused by auditory stimulation predict changes in memory consolidation.

## 4. Discussion

These findings demonstrate that auditory stimulation delivered during baseline activity or during the upstate of an endogenous SO during stages 2 and 3 NREM sleep reliably evokes SO and SO-spindle complexes activity in children with RE and control children. Evoked SOs and SO-spindle complexes are broadly distributed over the cortex, but the effect is maximal over the frontal electrodes near the midline. Frontal SO and SO-spindle complex rates positively predict sleep-dependent memory consolidation and changes in frontal SO and SO-spindle complex rates due to auditory stimulation positively predict changes in sleep-dependent memory consolidation. These results support the potential for closed-loop stimulation protocols that increase SO- and SO-spindle complex rates to enhance sleep-dependent memory consolidation.

Prior studies have demonstrated that auditory stimulation during stages 2 and 3 NREM sleep induces SOs and spindles^17-23^. Prior work has also suggested that auditory stimulation timed to the upstate of an endogenous SO is more effective at evoking SOs and spindles^17-20,22,46-48^. Here we strengthened these findings with improved resolution of the proportional impact of stimulation at different phases of the SO and also provide comparison to sham stimulation at the same phases to control for background activities. Taken together these findings confirm that phase-controlled auditory stimulation during sleep can evoke SOs and SO-spindle complex events. We also show that auditory stimulation evokes SO and SO-spindle complexes compared to sham when delivered on most background activity, except for during a SO downstate. Prior work has focused on the impact of auditory stimulation at the FZ or CZ electrodes^17-23,48^. Here, we extended this work to identify the spatial extent of the evoked signals. These findings suggest that SOs can be evoked over widespread cortical regions, but SO-spindle complexes are indeed maximally evoked over frontal electrodes near the midline. As some studies suggest that spindle activity in different cortical regions support domain specific memory consolidation^49-52^, this indicates that auditory stimulation may be more successful to enhance memory consolidation related to frontal lobe processing.

Although SO and SO-spindle complex event rates have been found to positively predict sleep dependent memory consolidation across multiple domains^22,26-28,53,54^, most prior work evaluating the impact of auditory stimulation did not compare memory performance to SO or SO-spindle complex rates^20,22,23,25^. When doing so, we found that the change in SO rate and SO-spindle complex event rate positively predicted the change in memory consolidation between the two visits. This is consistent with prior work indicating that sleep spindles in the frontal cortex predict memory consolidation of the motor sequence typing task^15,55,56^. Importantly, we found SO- and SO-spindle complexes are reliably evoked when stimulation is delivered during background activity in the absence of an endogenous slow oscillation or timed to the upstate of an endogenous slow oscillation. Taken together, these findings indicate that stimulations delivered frequently, both on background activity and also timed to the upstate of endogenous SO may be required to maximally increase SO- and SO-spindle complexes event rates.

Children with RE have deficits in memory consolidation relative to control children. We were underpowered to analyze our two groups separately, however we saw similar qualitative results in response to stimulation between children with RE and controls. Additionally, the unbalanced visit order in our cohort could have introduced order effects. To address this, we included visit order as a covariate in our statistical models, ensuring that the observed results were not confounded by visit order.

Together, these findings contribute evidence supporting the role of sleep oscillations in sleep-dependent memory consolidation, and insights on the impact of a non-invasive method to optimally control these oscillations in children with RE and typically developing controls. These results raise the possibility that tailored auditory stimulation approaches could serve as scalable, non-invasive therapeutic interventions to support memory consolidation and mitigate the memory impairments associated with epilepsy^15^.

## Acknowledgments

This work was supported by NINDS R01NS115868.

## References

1. Born J, Wilhelm I. System consolidation of memory during sleep. Psychol Res. Mar 2012;76(2):192–203. doi:10.1007/s00426-011-0335-6

2. Kwon H, Walsh KG, Berja ED, et al. Sleep spindles in the healthy brain from birth through 18 years. Sleep. Apr 12 2023;46(4)doi:10.1093/sleep/zsad017

3. Steriade M, Domich L, Oakson G, Deschenes M. The deafferented reticular thalamic nucleus generates spindle rhythmicity. J Neurophysiol. Jan 1987;57(1):260–73. doi:10.1152/jn.1987.57.1.260

4. McCormick DA, Bal T. Sleep and arousal: thalamocortical mechanisms. Annu Rev Neurosci. 1997;20:185–215. doi:10.1146/annurev.neuro.20.1.185

5. Schreiner T, Petzka M, Staudigl T, Staresina BP. Endogenous memory reactivation during sleep in humans is clocked by slow oscillation-spindle complexes. Nat Commun. May 25 2021;12(1):3112. doi:10.1038/s41467-021-23520-2

6. Steriade M, Nunez A, Amzica F. A novel slow (< 1 Hz) oscillation of neocortical neurons in vivo: depolarizing and hyperpolarizing components. The Journal of neuroscience : the official journal of the Society for Neuroscience. Aug 1993;13(8):3252–65. doi:10.1523/JNEUROSCI.13-08-03252.1993

7. Wodeyar A, Chinappen D, Mylonas D, et al. Thalamic epileptic spikes disrupt sleep spindles in patients with epileptic encephalopathy. Brain : a journal of neurology. Aug 1 2024;147(8):2803–2816. doi:10.1093/brain/awae119

8. Seibt J, Richard CJ, Sigl-Glockner J, et al. Cortical dendritic activity correlates with spindle-rich oscillations during sleep in rodents. Nat Commun. Sep 25 2017;8(1):684. doi:10.1038/s41467-017-00735-w

9. Buzsaki G. Hippocampal sharp wave-ripple: A cognitive biomarker for episodic memory and planning. Hippocampus. Oct 2015;25(10):1073–188. doi:10.1002/hipo.22488

10. Staresina BP, Bergmann TO, Bonnefond M, et al. Hierarchical nesting of slow oscillations, spindles and ripples in the human hippocampus during sleep. Nat Neurosci. Nov 2015;18(11):1679–1686. doi:10.1038/nn.4119

11. Dudai Y, Karni A, Born J. The Consolidation and Transformation of Memory. Neuron. 2015;88(1):20–32. doi:10.1016/j.neuron.2015.09.004

12. Geva-Sagiv M, Mankin EA, Eliashiv D, et al. Augmenting hippocampal–prefrontal neuronal synchrony during sleep enhances memory consolidation in humans. Nature Neuroscience. 2023/06/01 2023;26(6):1100–1110. doi:10.1038/s41593-023-01324-5

13. Ross EE, Stoyell SM, Kramer MA, Berg AT, Chu CJ. The natural history of seizures and neuropsychiatric symptoms in childhood epilepsy with centrotemporal spikes (CECTS). Epilepsy Behav. Feb 2020;103(Pt A):106437. doi:10.1016/j.yebeh.2019.07.038

14. Wickens S, Bowden SC, D’Souza W. Cognitive functioning in children with self-limited epilepsy with centrotemporal spikes: A systematic review and meta-analysis. Epilepsia. Oct 2017;58(10):1673–1685. doi:10.1111/epi.13865

15. Kwon H, Chinappen DM, Kinard EA, et al. Association of Sleep Spindle Rate With Memory Consolidation in Children With Rolandic Epilepsy. Neurology. 2025;104(2):e210232. doi:doi:10.1212/WNL.0000000000210232

16. Kramer MA, Stoyell SM, Chinappen D, et al. Focal Sleep Spindle Deficits Reveal Focal Thalamocortical Dysfunction and Predict Cognitive Deficits in Sleep Activated Developmental Epilepsy. The Journal of neuroscience : the official journal of the Society for Neuroscience. Feb 24 2021;41(8):1816–1829. doi:10.1523/JNEUROSCI.2009-20.2020

17. Ngo HV, Martinetz T, Born J, Molle M. Auditory closed-loop stimulation of the sleep slow oscillation enhances memory. Neuron. May 8 2013;78(3):545–53. doi:10.1016/j.neuron.2013.03.006

18. Ong JL, Lo JC, Chee NI, et al. Effects of phase-locked acoustic stimulation during a nap on EEG spectra and declarative memory consolidation. Sleep Med. Apr 2016;20:88–97. doi:10.1016/j.sleep.2015.10.016

19. Ngo HV, Seibold M, Boche DC, Molle M, Born J. Insights on auditory closed-loop stimulation targeting sleep spindles in slow oscillation up-states. Journal of neuroscience methods. Mar 15 2019;316:117–124. doi:10.1016/j.jneumeth.2018.09.006

20. Leminen MM, Virkkala J, Saure E, et al. Enhanced Memory Consolidation Via Automatic Sound Stimulation During Non-REM Sleep. Sleep. Mar 1 2017;40(3)doi:10.1093/sleep/zsx003

21. Besedovsky L, Ngo H-VV, Dimitrov S, Gassenmaier C, Lehmann R, Born J. Auditory closed-loop stimulation of EEG slow oscillations strengthens sleep and signs of its immune-supportive function. Nature Communications. 2017/12/07 2017;8(1):1984. doi:10.1038/s41467-017-02170-3

22. Baxter BS, Mylonas D, Kwok KS, et al. The effects of closed-loop auditory stimulation on sleep oscillatory dynamics in relation to motor procedural memory consolidation. Sleep. 2023;46(10)doi:10.1093/sleep/zsad206

23. Prehn-Kristensen A, Ngo HV, Lentfer L, et al. Acoustic closed-loop stimulation during sleep improves consolidation of reward-related memory information in healthy children but not in children with attention-deficit hyperactivity disorder. Sleep. Aug 12 2020;43(8)doi:10.1093/sleep/zsaa017

24. Bellesi M, Riedner BA, Garcia-Molina GN, Cirelli C, Tononi G. Enhancement of sleep slow waves: underlying mechanisms and practical consequences. Frontiers in systems neuroscience. 2014;8:208. doi:10.3389/fnsys.2014.00208

25. Harrington MO, Ngo HV, Cairney SA. No benefit of auditory closed-loop stimulation on memory for semantically-incongruent associations. Neurobiol Learn Mem. Sep 2021;183:107482. doi:10.1016/j.nlm.2021.107482

26. Hahn M, Joechner AK, Roell J, et al. Developmental changes of sleep spindles and their impact on sleep-dependent memory consolidation and general cognitive abilities: A longitudinal approach. Dev Sci. Jan 2019;22(1):e12706. doi:10.1111/desc.12706

27. Hahn MA, Heib D, Schabus M, Hoedlmoser K, Helfrich RF. Slow oscillation-spindle coupling predicts enhanced memory formation from childhood to adolescence. Elife. Jun 24 2020;9 doi:10.7554/eLife.53730

28. Astill RG, Piantoni G, Raymann RJ, et al. Sleep spindle and slow wave frequency reflect motor skill performance in primary school-age children. Frontiers in human neuroscience. 2014;8:910. doi:10.3389/fnhum.2014.00910

29. Mölle M, Born J. Slow oscillations orchestrating fast oscillations and memory consolidation. Prog Brain Res. 2011;193:93–110. doi:10.1016/b978-0-444-53839-0.00007-7

30. Klinzing JG, Tashiro L, Ruf S, Wolff M, Born J, Ngo HV. Auditory stimulation during sleep suppresses spike activity in benign epilepsy with centrotemporal spikes. Cell Rep Med. Nov 16 2021;2(11):100432. doi:10.1016/j.xcrm.2021.100432

31. Fattinger S, Heinzle BB, Ramantani G, Abela L, Schmitt B, Huber R. Closed-Loop Acoustic Stimulation During Sleep in Children With Epilepsy: A Hypothesis-Driven Novel Approach to Interact With Spike-Wave Activity and Pilot Data Assessing Feasibility. Hypothesis and Theory. Frontiers in human neuroscience. 2019-May-21 2019;13doi:10.3389/fnhum.2019.00166

32. Proposal for revised classification of epilepsies and epileptic syndromes. Commission on Classification and Terminology of the International League Against Epilepsy. Epilepsia. Jul-Aug 1989;30(4):389–99. doi:10.1111/j.1528-1157.1989.tb05316.x

33. Fisher RS, Acevedo C, Arzimanoglou A, et al. ILAE official report: a practical clinical definition of epilepsy. Epilepsia. Apr 2014;55(4):475–82. doi:10.1111/epi.12550

34. Wamsley EJ, Tucker MA, Shinn AK, et al. Reduced sleep spindles and spindle coherence in schizophrenia: mechanisms of impaired memory consolidation? Biological psychiatry. Jan 15 2012;71(2):154–61. doi:10.1016/j.biopsych.2011.08.008

35. Manoach DS, Pan JQ, Purcell SM, Stickgold R. Reduced Sleep Spindles in Schizophrenia: A Treatable Endophenotype That Links Risk Genes to Impaired Cognition? Biological psychiatry. Oct 15 2016;80(8):599–608. doi:10.1016/j.biopsych.2015.10.003

36. Manoach DS, Cain MS, Vangel MG, Khurana A, Goff DC, Stickgold R. A failure of sleep-dependent procedural learning in chronic, medicated schizophrenia. Biological psychiatry. Dec 15 2004;56(12):951–6. doi:10.1016/j.biopsych.2004.09.012

37. Grigg-Damberger M, Gozal D, Marcus CL, et al. The visual scoring of sleep and arousal in infants and children. J Clin Sleep Med. Mar 15 2007;3(2):201–40.

38. Kleiner M, Brainard DH, Pelli D, Ingling A, Murray R, Broussard C. What’s new in Psychtoolbox-3. Perception. 01/01 2007;36:1–16. doi:10.1068/v070821

39. Mölle M, Marshall L, Gais S, Born J. Grouping of spindle activity during slow oscillations in human non-rapid eye movement sleep. The Journal of neuroscience : the official journal of the Society for Neuroscience. Dec 15 2002;22(24):10941–7. doi:10.1523/jneurosci.22-24-10941.2002

40. Purcell SM, Manoach DS, Demanuele C, et al. Characterizing sleep spindles in 11,630 individuals from the National Sleep Research Resource. Nat Commun. Jun 26 2017;8:15930. doi:10.1038/ncomms15930

41. Iber C, Medicine AAoS. The AASM Manual for the Scoring of Sleep and Associated Events: Rules, Terminology and Technical Specifications. American Academy of Sleep Medicine; 2007.

42. Davis ZW, Muller L, Martinez-Trujillo J, Sejnowski T, Reynolds JH. Spontaneous travelling cortical waves gate perception in behaving primates. Nature. 2020/11/01 2020;587(7834):432–436. doi:10.1038/s41586-020-2802-y

43. Kwon H, Kronemer SI, Christison-Lagay KL, et al. Early cortical signals in visual stimulus detection. Neuroimage. Dec 1 2021;244:118608. doi:10.1016/j.neuroimage.2021.118608

44. Oostenveld R, Fries P, Maris E, Schoffelen JM. FieldTrip: Open source software for advanced analysis of MEG, EEG, and invasive electrophysiological data. Comput Intell Neurosci. 2011;2011:156869. doi:10.1155/2011/156869

45. Nishida M, Walker MP. Daytime naps, motor memory consolidation and regionally specific sleep spindles. PloS one. Apr 4 2007;2(4):e341. doi:10.1371/journal.pone.0000341

46. Batterink LJ, Creery JD, Paller KA. Phase of Spontaneous Slow Oscillations during Sleep Influences Memory-Related Processing of Auditory Cues. The Journal of neuroscience : the official journal of the Society for Neuroscience. Jan 27 2016;36(4):1401–9. doi:10.1523/jneurosci.3175-15.2016

47. Cox R, Korjoukov I, de Boer M, Talamini LM. Sound asleep: processing and retention of slow oscillation phase-targeted stimuli. PloS one. 2014;9(7):e101567. doi:10.1371/journal.pone.0101567

48. Henin S, Borges H, Shankar A, et al. Closed-Loop Acoustic Stimulation Enhances Sleep Oscillations But Not Memory Performance. eneuro. 2019;6(6):ENEURO.0306-19.2019. doi:10.1523/eneuro.0306-19.2019

49. Schmidt C, Peigneux P, Muto V, et al. Encoding difficulty promotes postlearning changes in sleep spindle activity during napping. The Journal of neuroscience : the official journal of the Society for Neuroscience. Aug 30 2006;26(35):8976–82. doi:10.1523/JNEUROSCI.2464-06.2006

50. Gais S, Molle M, Helms K, Born J. Learning-dependent increases in sleep spindle density. The Journal of neuroscience : the official journal of the Society for Neuroscience. Aug 1 2002;22(15):6830–4. doi:20026697

51. Fernandez LMJ, Lüthi A. Sleep Spindles: Mechanisms and Functions. Physiological Reviews. 2020;100(2):805–868. doi:10.1152/physrev.00042.2018

52. Morin A, Doyon J, Dostie V, et al. Motor sequence learning increases sleep spindles and fast frequencies in post-training sleep. Sleep. Aug 2008;31(8):1149–56.

53. Muehlroth BE, Sander MC, Fandakova Y, et al. Precise Slow Oscillation–Spindle Coupling Promotes Memory Consolidation in Younger and Older Adults. Scientific reports. 2019/02/13 2019;9(1):1940. doi:10.1038/s41598-018-36557-z

54. Ng T, Noh E, Spencer RMC. Does slow oscillation-spindle coupling contribute to sleep-dependent memory consolidation? A Bayesian meta-analysis. eLife Sciences Publications, Ltd; 2024.

55. Jacobacci F, Armony JL, Yeffal A, et al. Rapid hippocampal plasticity supports motor sequence learning. Proceedings of the National Academy of Sciences. 2020;117(38):23898–23903. doi:doi:10.1073/pnas.2009576117

56. Spencer ER, Chinappen D, Emerton BC, et al. Source EEG reveals that Rolandic epilepsy is a regional epileptic encephalopathy. NeuroImage Clinical. 2022;33:102956. doi:10.1016/j.nicl.2022.102956

